# Percutaneous Intramyocardial Septal Cryoablation for Hypertrophic Cardiomyopathy: A Canine Model with 6 months Follow-up

**DOI:** 10.1101/2023.08.08.552550

**Authors:** Xiaonan Lu, Jing Li, David H. Hsi, Juan Zhang, Yupeng Han, Shengjun Ta, Jing Wang, Jing He, Jia Zhao, Chao Hao, Lu Yao, Xumei Ou, Bo Shan, Bo Wang, Xueli Zhao, Rui Hu, Liwen Liu

## Abstract

**BACKGROUND:** Echocardiography-guided percutaneous intramyocardial septal radiofrequency ablation (PIMSRA, Liwen procedure) is a novel treatment option for hypertrophic obstructive cardiomyopathy (HOCM). The safety and feasibility of using this procedure for cryoablation are unknown.

**OBJECTIVE:** To establish a canine model for echocardiography-guided percutaneous intramyocardial septal cryoablation (PIMSCA).

**METHODS:** Eight canines underwent PIMSCA and had electrocardiography, echocardiography, myocardial contrast echocardiography (MCE), (ECG), serological and pathological examinations during the preoperative, immediate postoperative, and in 6-month follow-up.

**RESULTS:** All eight canines underwent successful cryoablation and continued to be in sinus rhythm during ablation and without malignant arrhythmias. MCE showed that ablation area had decreased myocardial perfusion after procedure. Troponin I levels were significantly elevated [0.010 (0.005, 0.297) ng/mL vs. 3.122 (1.152,7.990) ng/mL, p < 0.05)]. At follow-up 6-months after procedure, all animals were alive, with thinning of the interventricular septum (7.26 ± 0.52 mm vs. 3.86 ± 0.29 mm, p < 0.05). Echocardiography showed no significant decrease in left ventricular ejection fractions (LVEF) (54.32 ± 2.93 %vs. 54.70 ± 2.47 %, p > 0.05) or changes by pulse-wave Doppler E/A (1.17 ± 0.43 vs. 1.07 ± 0.43, p > 0.05), E/e’(8.09 ± 1.49 vs. 10.05 ±2.68, p > 0.05). Pathological findings proved effective cryoablation in myocardial tissues. We observed pericardial effusions and premature ventricular complexes (PVCs) associated with the procedure.

**CONCLUSION:** Our findings provide preliminary evidence of the safety and feasibility of PIMSACA in interventricular septum reduction, which provides a potentially new treatment option for HOCM.

## Introduction

Hypertrophic cardiomyopathy (HCM) is the most common inherited cardiovascular disease and one of the most common causes of sudden cardiac death (SCD) in young adolescents and athletes^1 2^. Approximately two-thirds of patients have left ventricular outflow tract (LVOT) obstruction known as hypertrophic obstructive cardiomyopathy (HOCM), which may result in poor prognosis ^3^. We published our data on transthoracic echocardiography–guided percutaneous intramyocardial septal radiofrequency ablation (PIMSRA, Liwen procedure) for HOCM. The safety and feasibility of PIMSRA have been confirmed in our previous reports ^4–6^.

PIMSRA (Liwen procedure) is based on radiofrequency ablation (RFA). RFA works by heating the tissue to hyperthermic levels for several minutes resulting in necrosis without a sharp boundary to the ablation zone ^7^, It is technically demanding for the operator, and requires accumulated experience to perfect operator skills.Cryoablation causes tissue injury through a process of freezing and rewarming. The temperatures of local tissues rapidly change, and cryogenic damage occurs, creating tissue necrosis with a clear margin.

Consequently, we hypothesized using percutaneous intramyocardial cryoablation could obtain the same ablation effect and provide a new option for HOCM treatment. We designed and implemented a canine model to evaluate the safety and feasibility of PIMSCA for the treatment of HOCM.

## Materials and Methods

### Preparation

Eight canines were raised in the Laboratory Animal Center of Xijing Hospital. This study complied with the Helsinki Declaration and was approved by the ethics reviews committee of Xijing Hospital (KY20162042-1). All canines were clinically determined to be in good health, fasted for 24 hours before the procedures. The age of canines was 22-24 months, mean weight was 17±5.04 kg, and 50% were male. Before the procedure, they were anesthetized with 3% sodium pentobarbital (1 ml/kg) injected by the hindlimb vein. Transthoracic echocardiography (TTE) (EPIQ 7C, Philips Ultrasound Inc., USA), standard 6-lead electrocardiography (ECG), blood biochemical, and myocardial contrast echocardiography (MCE) examination were performed. Myocardial biopsy was achieved by echo guided percutaneous intramyocardial septal pathway ^8^ to obtain the myocardial tissue from the interventricular septum.

TTE was performed on an EPIQ 7C ultrasound instrument equipped with a 1- to 5-MHz carrier frequency transducer. We measured the following parameters :

- Thickness of the septum
- M-mode Amplitude of movement of the ablated region
- Left ventricular ejection fraction (LVEF)
- Stroke volume (SV)
- Stroke volume index (SVI).
- M-mode echo Amplitude of the motion of the regional interventricular septum
- Ratio of peak early mitral inflow velocity (E-wave) and peak late mitral inflow velocity (A-wave), E/A ratio on the pulsed-wave (PW) Doppler,
- Ratio of early diastolic trans-mitral flow velocity (E) to tissue Doppler early-diastolic septal mitral annular velocity (septal E/e′).

Subsequently, myocardial contrast echocardiography (MCE) was performed using continuous infusion of Sonovue® (Bracco Imaging) to evaluate myocardial perfusion. All echocardiographic examinations were performed by a trained sonographer in compliance with American Society of Echocardiography (ASE) guidelines ^9^.

### PIMSCA procedure

The procedure process is illustrated in Fig 1. Under the guidance of TTE, the cryoprobe (15G, Endocare Cryocare System, USA) was inserted into the interventricular septum to avoid the surface of the coronary blood vessels, then advanced to the basal segment of the interventricular septum (ensuring the distance of 8-10 mm between the cryoprobe tip and aortic valve to avoid the membranous segment). PIMSCA was started at 100% cryoablation power for 90 seconds. On echocardiography, the cryoablation area was demonstrated by arc hypoechoic with acoustic shadows (Fig 1, AI-II). When thawed to 10°C, the cryoprobe was rapidly withdrawn, and the puncture site was pressed briefly with a sterile dry cotton ball. Simultaneous ECG recordings were monitored (1550P, Nihon Kohden, Japan) constantly during the procedure.

**Fig.1.**
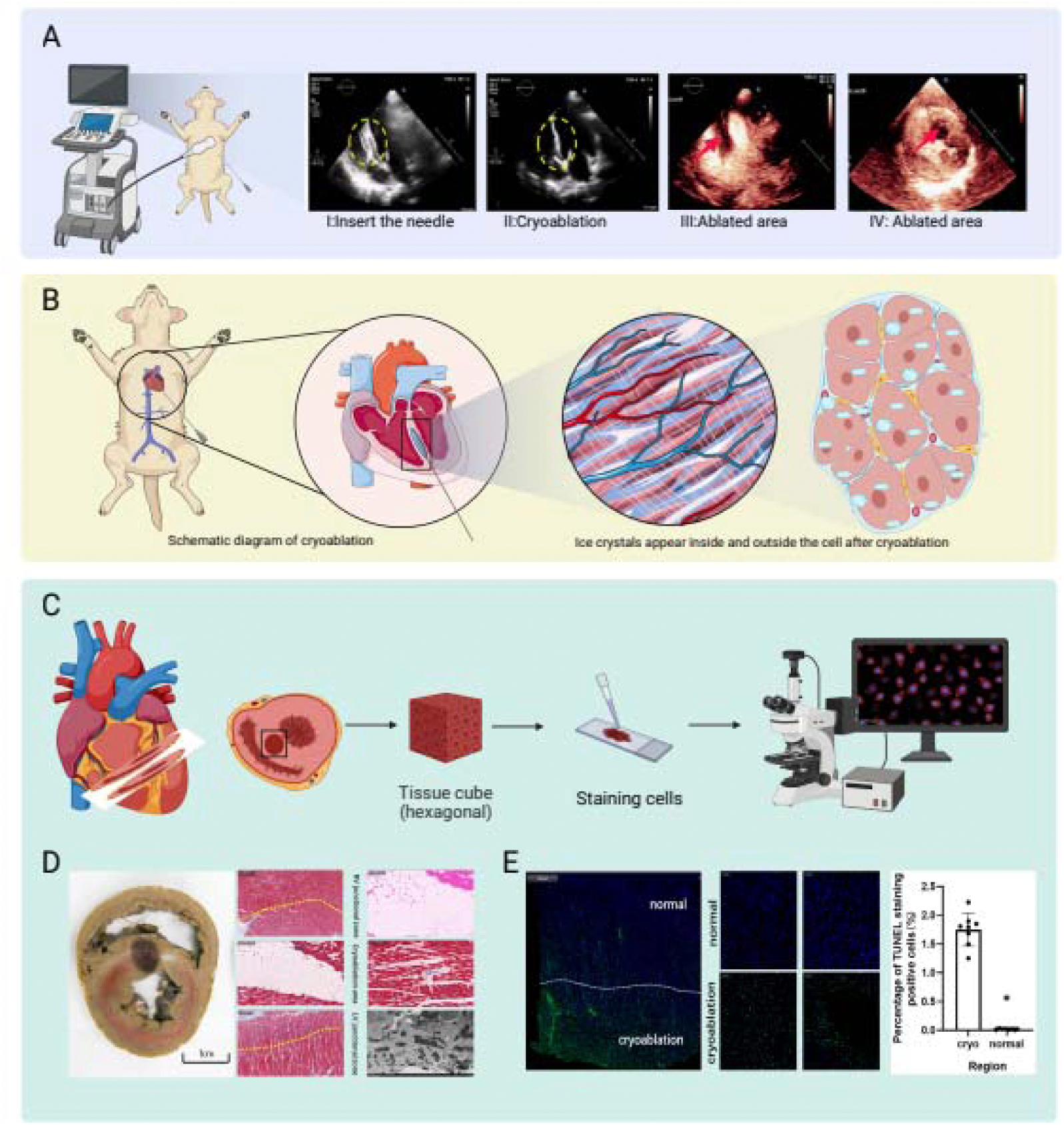
Echocardiography-guided percutaneous intramyocardial septal cryoablation: A: I: Insert the needle, II: After cryoablation, a curved strong echo accompanied by sound shadow appeared. III-IV: Ablation area displayed by MCE after cryoablation (Black void). B: The illustration of echocardiography-guided percutaneous intramyocardial septal cryoablation (PIMSCA). C: Immediate postoperative tissueprocessing. D: After 48hour fixation with paraformaldehyde cardiac short-axis section specimens, necrotic areas shown by Masson staining (×40), HE staining (×40), and electron microscope observation (×4200). E: The percentage of TUNEL stained (×20) cells immediately after cryoablation. Eight pictures were selected for analysis in each field, and the percentage of TUNEL positive cells versus normal myocardium in the cryoablation area was determined. Created with BioRender.com, parts of the figure have been created using SMART – Servier Medical Art, Servier: https://smart.servier.com.

Perfusion defects in the ablated regions were assessed by MCE after the procedure and compared to this area preoperatively to verify successful cryoablation. We took samples at the cryoablation region and adjacent regions. Following harvesting, the specimens were fixed with 2% paraformaldehyde and 2.5% glutaraldehyde for hematoxylin-eosin (HE) staining, Masson staining, TUNEL staining, and transmission electron microscopy (TEM) examination (HT-7800, Hitachi, Japan). The ablated area (8 images per ablated area) was analyzed using (Image-Pro 7.0, NIH, USA.) Cryoablation effects were assessed macroscopically and microscopically.

### Follow-up

Immediately after the procedure we performed TTE, 6-lead ECG, and reviewed them 1, 3, and 6 months after the procedure. We performed the blood biochemical analysis one hour after the procedure. A pathological examination was performed at 6 months postoperatively in order to verify the effectiveness of ablation. Follow-up outcomes included safety and feasibility indicators. Primary endpoints included morbidity and mortality; and secondary endpoints were pericardial effusion, premature ventricular contractions. Feasibility hypothesis was that cryoablation could produce effective necrosis with thinning of the ablation zone at the end of follow-up.

### Statistical analysis

Statistical analyses were performed using SPSS (SPSS 17.0, IBM SPSS Statistics, USA). We first assessed whether the data conformed to the normal distribution. Quantitative variables were expressed as means ± standard deviation (SD) or median (quartile 1, quartile 3). Paired comparisons were performed using Wilcoxon signed-rank tests as appropriate. we used linear mixed effects models for repeated measures was used to compare continuous variables over time, Bonferroni correction for multiple comparisons were used for pairwise comparisons. The significance level was set at 0.05, and P values less than 0.05 indicated a statistically significant difference.

## Results

### Baseline characteristics of the canines

Eight canines were of age 22-24 months, and weighed 17±5.04 kg, the ventricular septal thickness was 7.26 ± 0.52mm (Table1). Preoperative myocardial biopsy results are shown in Figure 3. The myocardial cells are arranged neatly with uniform size and shapes. Immunofluorescence staining of TUNEL revealed that DNA fragmentation was intact (Fig.1). Transmission electron microscopy (TEM) revealed orderly arrangement of myofilaments, discs neat and continuous, normal shape and structure of mitochondria, and an orderly arrangement and clear structure of cristae in canine cardiomyocytes (Fig.3).

**Table1:**
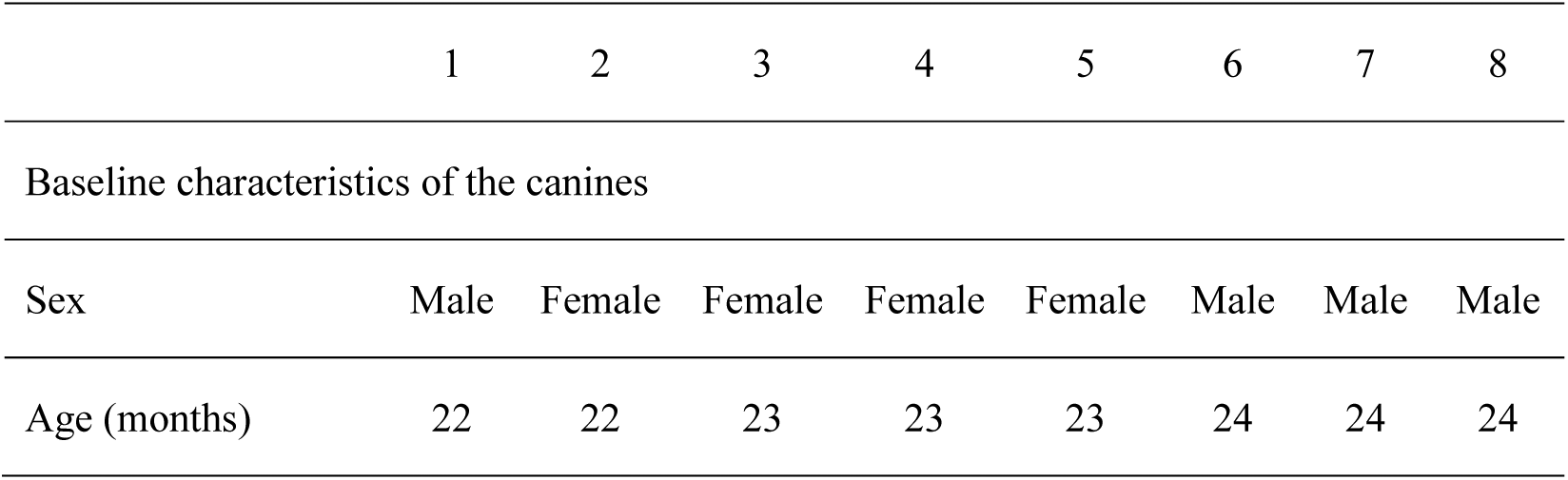
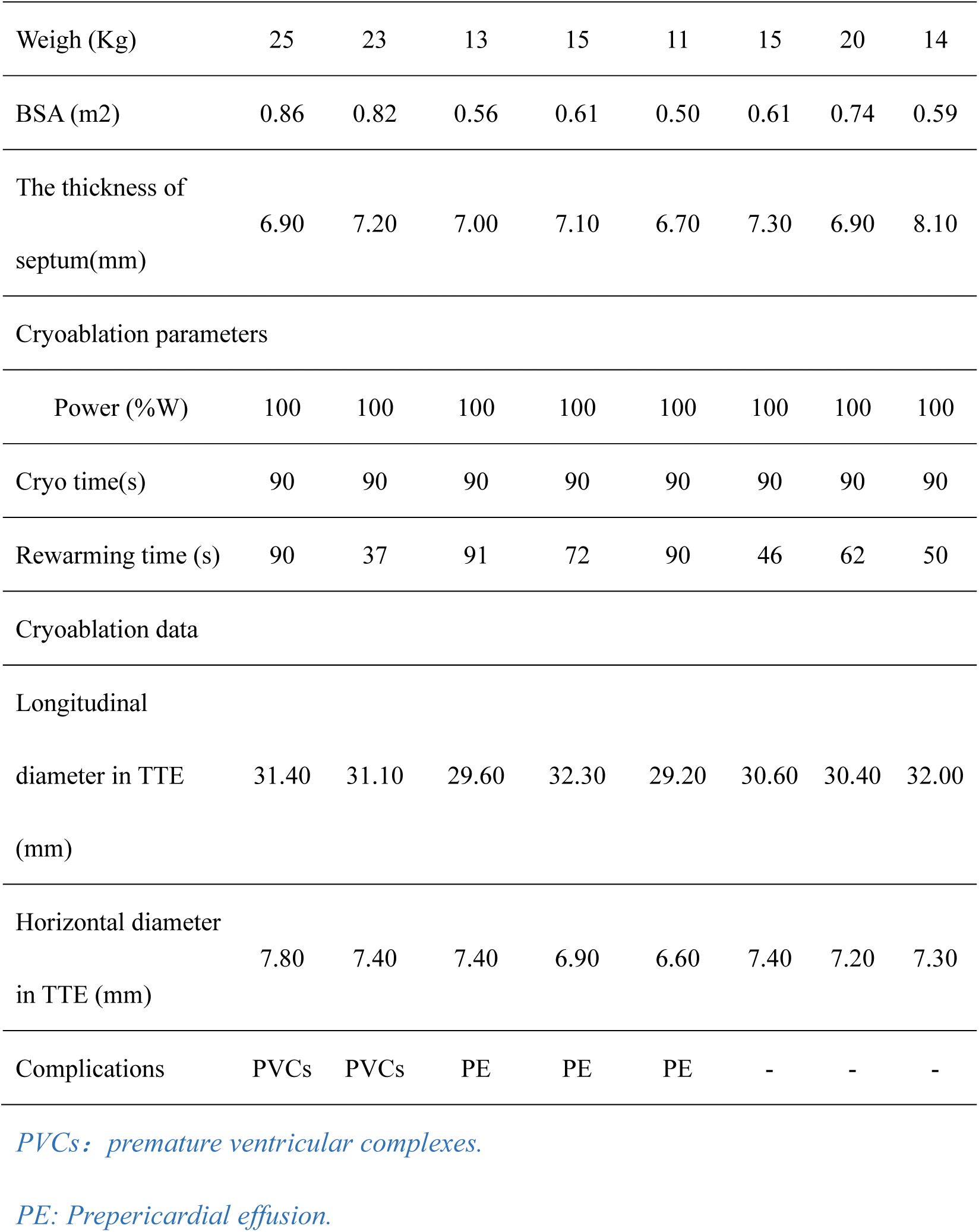
Baseline information.

### Safety of PIMSCA

All of the canines survived the instrumentation and the experiment without complications such as pericardial tamponade or ventricular fibrillation). No arrhythmia developed during cryoablation. Heart rate of the canines were of no significant changes from the baseline during procedure (Fig. 2) (121.5 ± 12.49 vs. 123.38 ± 24.81, p > 0.05). Canines survived to 6-month follow-up. In addition, no evidence for bundle branch block or other abnormalities were observed on the ECG (Fig. 1). The number 7 canines had a pulmonary infection triggered by injury to the left hind paw, and was excluded to ensure the accuracy and authenticity of the data.

**Fig.2.**
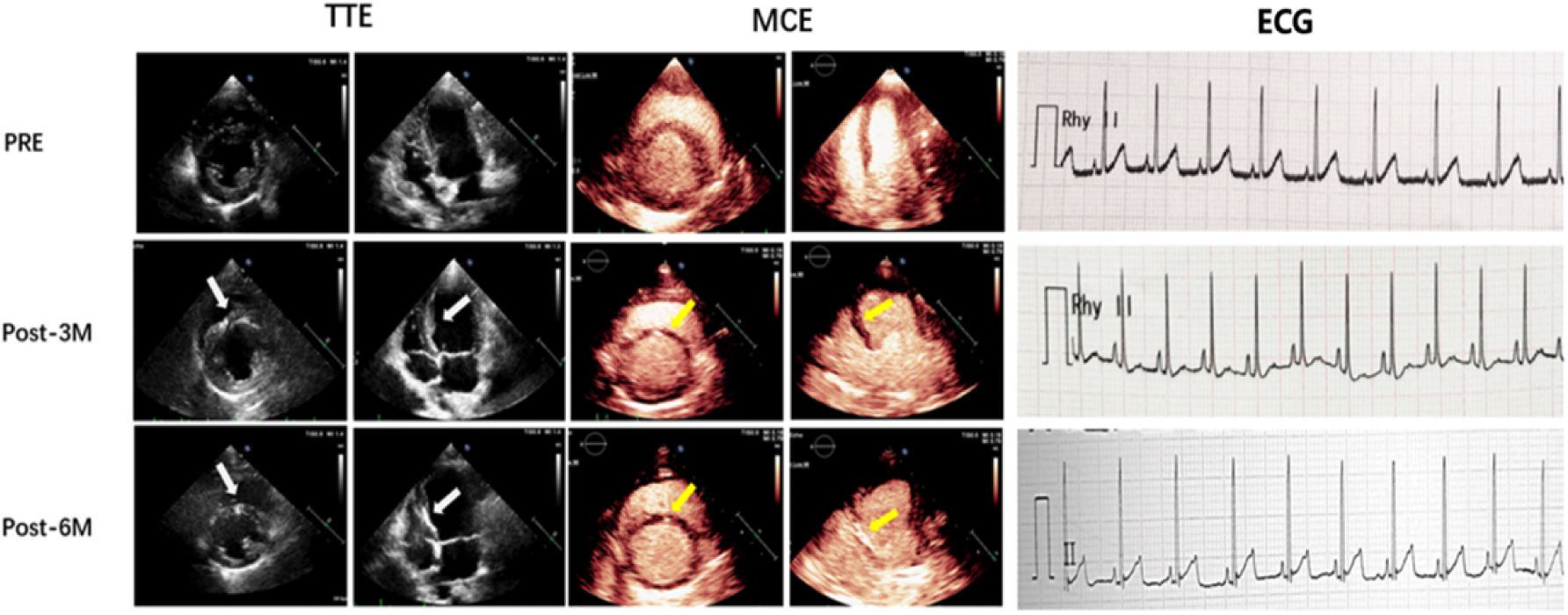
Echo, contrast echo and lead II electrocardiogram before the procedure, 3-month, and 6-month after the procedure.

**Fig.3.**
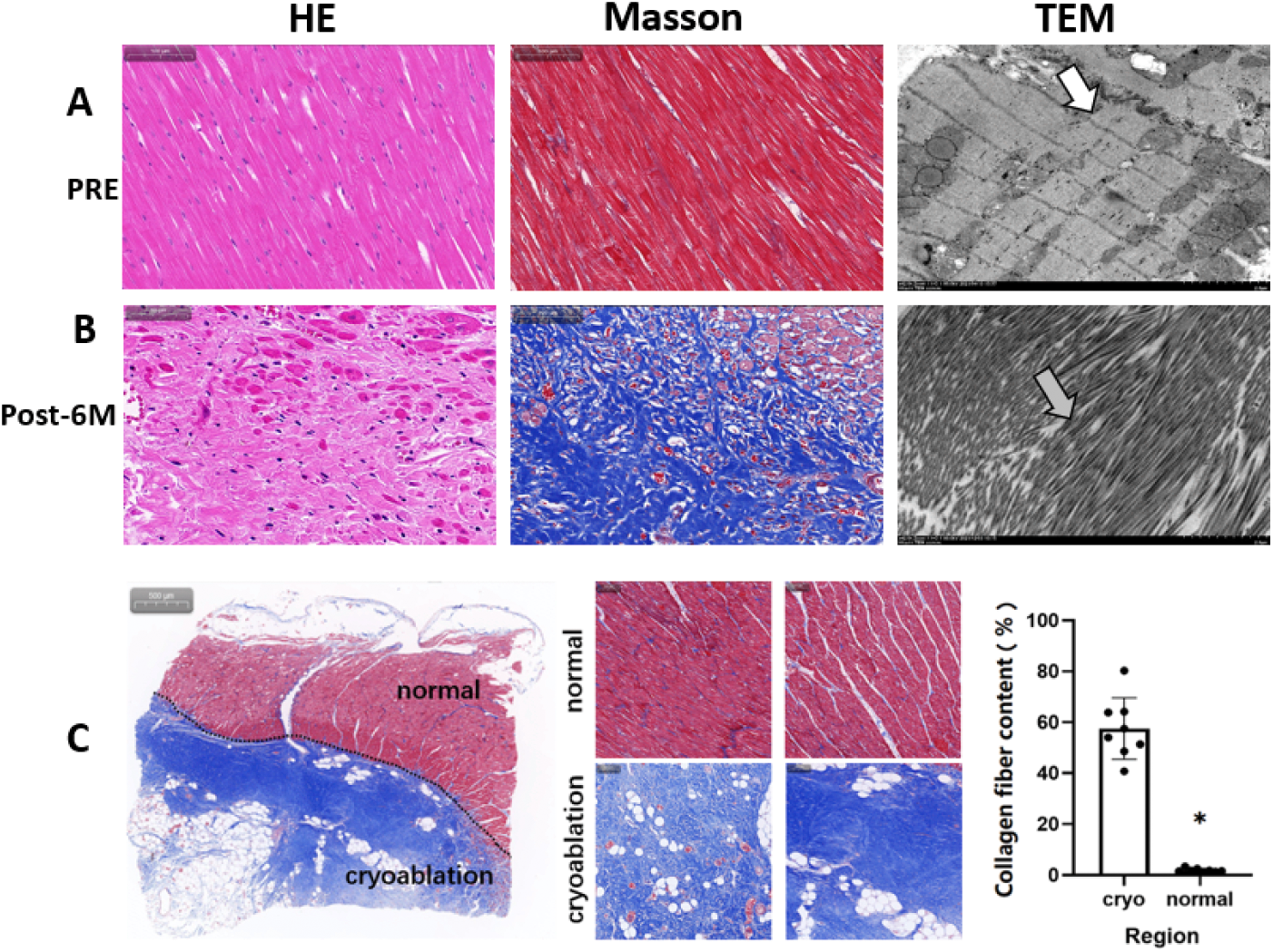
HE staining (×40), Masson staining (×40), TUNEL staining (C, G, K) (×20) and electron microscope observation (×4200) (D, H, L) of the cryoablation interventricular septum. Mitochondria (hollow arrow), myofibril (grey arrow). HE: hematoxylin-eosin staining, TEM: Transmission electron microscopy. *p < 0.05.

For a detailed analysis of the variation in the thickness and cardiac function during the 6-month follow-up, we performed echo to quantify the global and regional cardiac functions (Table1). Compared to the pre-operative level, echocardiography showed no significant change in Left ventricular ejection fractions (LVEF) (54.32 ± 2.93 vs. 54.70 ± 2.47 %, p > 0.05) and E/A (1.17 ± 0.43 vs. 1.07 ± 0.43, p > 0.05), E/e’(8.09 ± 1.49 vs. 10.05 ±2.68, p > 0.05). After the operation, the value of stroke volume index (SVI) of the left ventricle and the longitudinal strain of the ablated interventricular septum region were reduced, and these parameters gradually returned to the pre-operative level during the 6-months.

### Feasibility of PIMSCA

Fig. 2 and Fig.4 shows the variation in the interventricular septum on TTE and MCE follow-up after the procedure, progressive thinning of septal thickness on echocardiography over time, the thickness of the ablated interventricular septum significantly decreased from 7.26 ± 0.52 to 3.86 ± 0.29 mm (p < 0.05) at 6-months after the procedure. Six months after the procedure, gradual absorption of perfusion defects on MCE showed enhancement compared with the surrounding myocardium (Fig.2) Hypoperfusion can be seen in the interventricular septal area. New capillary formation after myocardial necrosis was observed.

**Fig.4.**
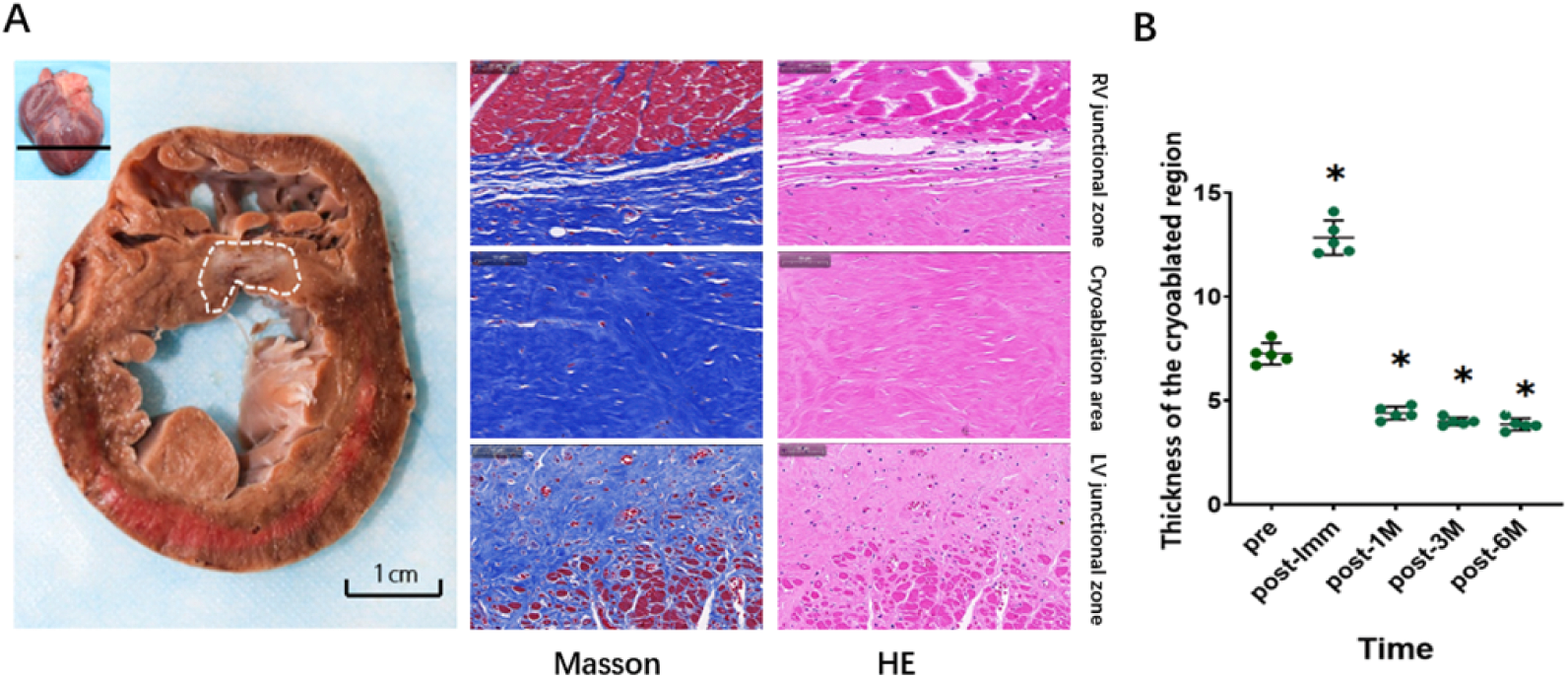
A: Pathological changes in the ablation area 6-months after the procedure General view of the heart. B: Changes in interventricular septal thickness. * p < 0.05 vs. before PIMSCA.

Myocardial cells appeared swollen and exhibited irregular morphologies, and the gap between cells increased on HE staining. Masson staining showed the collagen and myocardial fibers were broken. TUNEL staining (Central Illustration) demonstrated DNA fragmentation in cryoablation area more than normal area (1.75 ± 0.29 versus, 0.02 ± 0.01%; P < 0.000). Mitochondrial swelling, vacuolation of some mitochondria, disappearance of mitochondrial cristae, dilatation of the sarcoplasmic reticulum, red blood cell leakage, disordered arrangement of myofilaments and sarcomeres, and myofilament disruption were observed under electron microscopy (Central Illustration). The Troponin I level significantly higher than the pre-operative level [0.010 (0.005, 0.297) ng/mL vs. 3.122 (1.152,7.990) ng/mL, p < 0.05)].

Six months after the procedure, cryoablation of the necrotic area was absorbed and remodeled, leaving a small, fixed fibrous scar (Fig.4, Irregular white dashed coil) was located in the mid interventricular septum. The area and macroscopically appeared white and significantly thinner than before the procedure and in other parts of the same septum. HE staining and Masson staining showed clear ablation boundaries; and the tissue in the ablation area was replaced by fibrous tissues (57.58±12.04 versus 1.89±0.71%; P<0.000, Fig.4). No normal cells were found by TUNEL staining. Under a magnification of ×4200 on electron microscope, the details after ablation and neovascularization was visible in collagen fibers.

### Complications

Three canines had pericardial effusions; two of which required percutaneous drainage without further problems; the other one did not require intervention. Pericardial effusion was not observed at one-week follow-up. Because Canine’s heart is smaller than human’s, 1.7 mm diameter cryo-needle tip probably injured blood vessels during the entrance into the heart. We believe that in humans with truly hypertrophic interventricular septum, echo-guided cryoprobe insertion can avoid blood vessels and significantly reduce the incidence of pericardial effusion using pre-operative computer tomography and intra-op echo imaging guidance as described in our previous publications ^4,6^. During cryoprobe insertion, two canines had transient premature ventricular complexes (PVCs), which disappeared after minor needle adjustment.

The amplitude of the motion (Table 2) of the interventricular septum decreased significantly on the M-mode (5.26 ± 0.05 vs. 2.17 ± 0.04, p > 0.05) after 6-months, which indicated effective septum reduction procedure.

**Table2:**
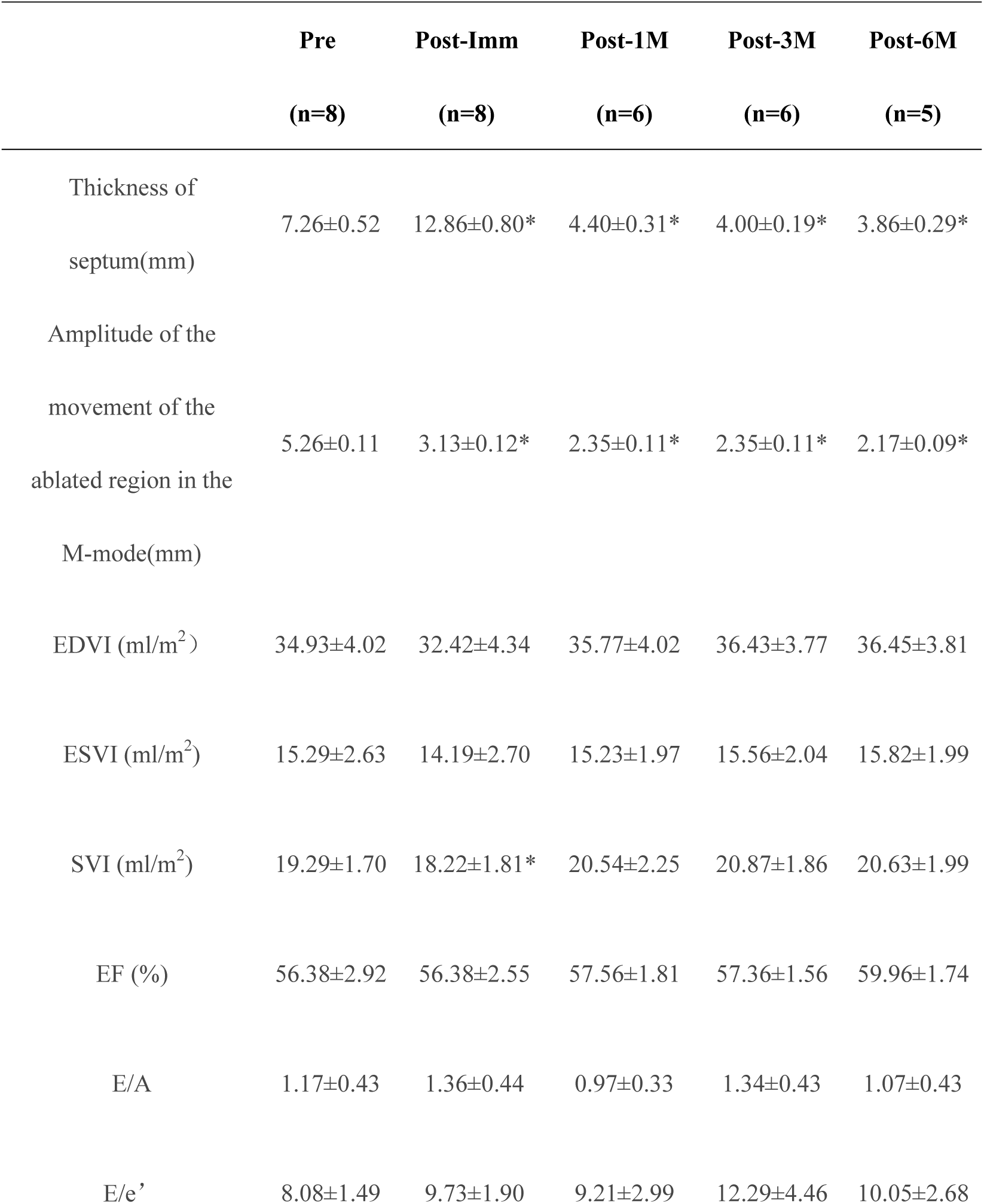
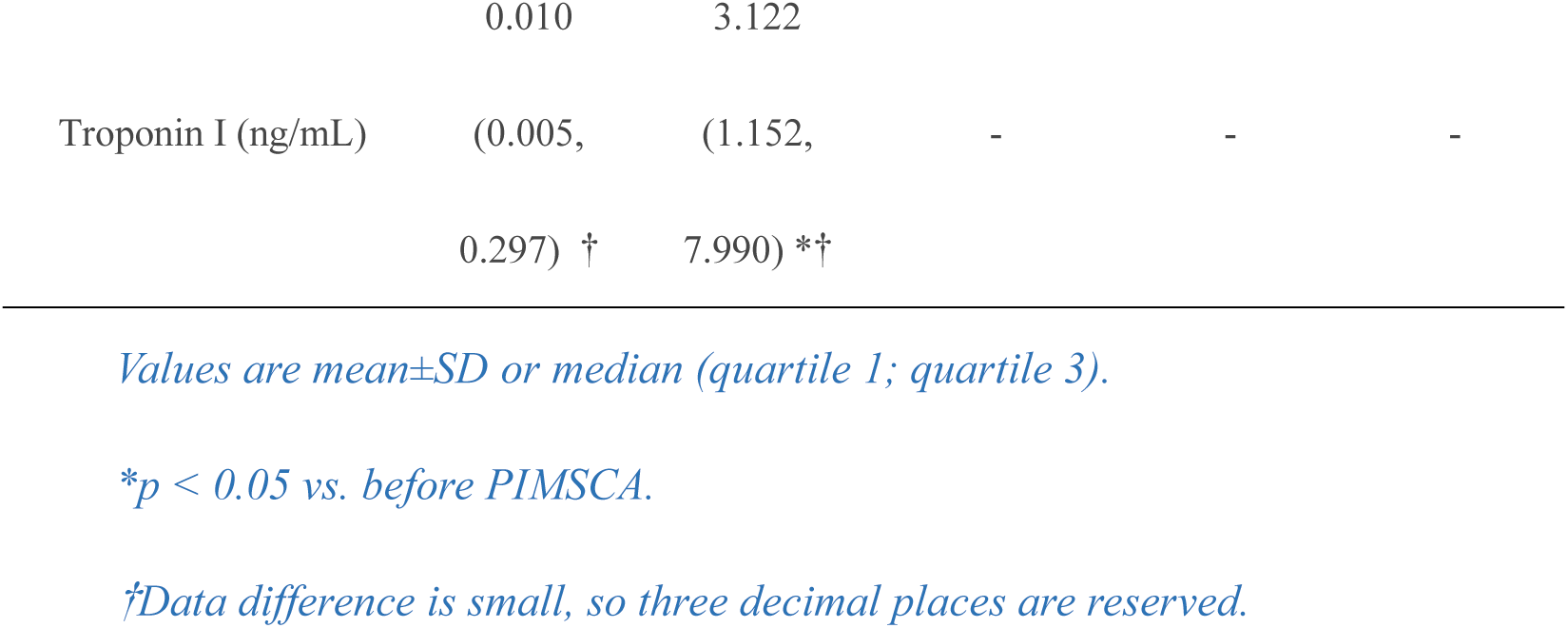
Comparison of experimental values in the preoperative, immediate postoperative and follow-up phases of PIMSCA (x̅±s)

## Discussion

PIMSCA (Liwen procedure) is a promising treatment option in HOCM. Our study focused on the short-term response of echo-guided percutaneous cryo-ablation of the healthy canine ventricular septum. All canines survived PIMSCA with thinning of the interventricular septum at all visits for 6-months postoperatively. Thus, PIMSCA appeared technically feasible and fairly safe; and provides a possibly new treatment option for HOCM patients.

In this study, we performed a novel procedure in the beating heart, via direct apical access to the interventricular septum. We used Echo and ECG to monitor cryoablation area in real-time. After the procedure, myocardial contraction could occlude small puncture sites after cryoprobe extraction reducing the risk of intraoperative bleeding and arrhythmia.

No arrhythmias, such as left bundle branch block (LBBB), or right bundle branch block (RBBB) occurred during the procedure and 6-months follow-up of cryoablation. The borders of the cryoablation were well defined, using echocardiography and ECG monitoring of the ablation zone in real-time. Cryo-ablation was stopped immediately if arrhythmias were noted. The cryoablation zone was 10 mm from the aortic valve annulus to avoid membranous septum and conduction delays. The left and right bundle branches are located under the endothelium of the ventricular septum ^8^. Our ablation injury is located within the myocardium, with a safe distance of 3-5 mm between the ablation needle tip and the endocardial surface, and minimizing endocardial injury to the intraventricular conduction system in the sub-endocardium). The short period of hypothermia in the tissue may cause reversible injury to the conduction bundle if occurred ^10^. Our study made it feasible to establish precise ablation in healthy canines with an interventricular septum thickness of 6-8 mm. In patients with HOCM, the average interventricular septum thickness is typically > 20 mm ^11^, which should leave enough room for safe ablation and avoid injury to the conduction system.

Left ventricular stroke volume index (SVI) decreased immediately after surgery and returned to normal in 1-month after the operation, which we believe is related to the postoperative tissue edema. We noticed that stroke volume index (SVI) recovery time is approximately the same as the time for edema to subside after cryoablation. We believe that PIMSCA is a safe and effective method for septum reduction, with good and stable efficacy.

Pericardial effusion was observed during PIMSCA procedure, which required pericardiocentesis but no persistent effusion after the procedure. We believe that in humans with truly hypertrophic interventricular septum, echo-guided cryoprobe insertion can avoid blood vessels and significantly reduce the incidence of pericardial effusion using pre-operative computer tomography imaging and intra-op echo guidance as described in our previous publications^4,6^. Cryoablation applied freezing and rewarming processes produce injuries to the cells, causing ice crystal formation and cell death ^12^. Freezing damage to the endothelium results in increased capillary permeability, causing bleeding, edema, and induced platelet aggregation, which forms microthrombus ^13^. Apoptosis increased in the sub-cold zone around the cryoablation area ^14^.

Immediately after the procedure (Fig.1), HE staining showed cells swollen and exhibiting irregular morphologies, and the gap between cells increased. Masson staining showed that the collagen and myocardial fibers were broken. TUNEL staining demonstrated DNA fragmentation in cryoablation area. Mitochondrial swelling, vacuolation of some mitochondria, disappearance of mitochondrial cristae, dilatation of the sarcoplasmic reticulum, red blood cell leakage, and disordered arrangement of myofilaments and sarcomeres, as well as myofilament disruption, were observed under electron microscopy. By comparing myocardial pathological alterations, we found that effective myocardial injury was present in the cryoablation area. Troponin I level was significantly higher than the pre-operative level. Troponin I was a myocardial regulatory protein showing high sensitivity and specificity in myocardial injury or necrosis ^15,16^, which also increased after alcohol ablation ^17^ or radiofrequency ablation ^18^, which was evidence of effective myocardial injury induced by cryoablation energy.

At 6 month follow up, the septal thickness was significantly thinned after the procedure; cryoablation of the necrotic area was absorbed and remodeled, leaving a small, fixed fibrous scar located in the mid interventricular septum, validating the effectiveness of cryoablation.

### Limitations

In this study, the peak LVOT gradient before and after the procedure was ≤3 mmHg, because of the healthy animal model. The most commonly established animal models of HCM are limited to small animal models ^19–22^(i.e., mice, rats, rabbits, felines), These animals are too small for myocardial ablation procedures. The ablation dose in these animals is far from being a meaningful parameter for clinical requirements ^23^. The development of rhythm disturbances after cryo-ablation may require prolonged ECG monitoring such as Holter ECG) and cannot be assessed based on snapshot ECGs. Finally, our canine model research had very small subject numbers due to ethics and resource considerations.

## Conclusion

The PIMSCA canine model is technically feasible and safe without mortality. We presented biochemical, echo, and pathology results of cryo-ablation. We can reasonably estimate that cryoablation is effective in septum reduction. This study highlights a new approach to reduce septal tissue and provides supporting evidence for the use of cryoablation in future clinical trials of HOCM treatment.

## Ethics approval

The Institutional Ethics Committee (KY20162042-1) of Xijing Hospital approved the procedure, which was performed in accordance with the ethical standards of the Declaration of Helsinki.

